# Epithelial WNT2B and Desert Hedgehog are necessary for human colonoid regeneration after bacterial cytotoxin injury

**DOI:** 10.1101/434639

**Authors:** Julie G. In, Jianyi Yin, Michele Doucet, Robert N. Cole, Lauren DeVine, Mark Donowitz, Nicholas C. Zachos, Sarah E. Blutt, Mary K. Estes, Olga Kovbasnjuk

## Abstract

Intestinal regeneration and crypt hyperplasia after radiation or pathogen injury relies on Wnt signaling to stimulate stem cell proliferation. Mesenchymal Wnts are essential for homeostasis and regeneration in mice, but the role of epithelial Wnts remains largely uncharacterized. Using the enterohemorrhagic *E. coli* secreted cytotoxin, EspP to induce injury to human colonoids, we evaluated a simplified, epithelial regeneration model that lacks mesenchymal Wnts. Here, we demonstrate that epithelial-produced WNT2B is upregulated following injury and essential for regeneration. Hedgehog signaling, specifically activation via the ligand Desert Hedgehog (DHH), but not Indian or Sonic Hedgehog, is another driver of regeneration and modulates WNT2B expression. These findings highlight the importance of epithelial WNT2B and DHH in regulating human colonic regeneration after injury.

## INTRODUCTION

The adult intestine has the amazing capacity to regenerate following stress, inflammation, or injury (Beumer and Clevers, 2016); however, the mechanisms that regulate regeneration are not well understood. Much of our knowledge in intestinal stem cell renewal and regeneration stems from studies in *Drosophila* (Jiang et al., 2016) and mice (Farin et al., 2016; Metcalfe et al., 2014; Ritsma et al., 2014). Studies in mouse models have led to characterization of the active and reserve intestinal stem cells in homeostasis and injury. Particularly relevant are *Drosophila* studies that revealed the importance of Wnt and Hedgehog signaling in development, maintenance, and regeneration of the midgut. However, the interplay of these two signaling pathways is not limited to intestinal maintenance. Hedgehog and Wnt signaling are essential pathways in development, homeostasis, and regeneration of many organs. The common features that influence regeneration after injury in classical regeneration models are: Wnt, Hedgehog, and Notch (Franco et al., 2013). Hedgehog signaling is essential in skin wound healing (Le et al., 2008), cardiac (Wang et al., 2016a), gastric (Konstantinou et al., 2016), lung (Sriperumbudur et al., 2016), hematopoietic (Trowbridge et al., 2006), and liver regeneration (Langiewicz et al., 2016; Wang et al., 2016b), as well as epidermal stem cell homeostasis (Adolphe, 2004). Additionally, intestinal regeneration in *Drosophila* is stimulated by active Hedgehog signaling (Tian et al., 2015). *Sonic hedgehog (SHH)* is the most widely expressed mammalian Hedgehog ligand (Varjosalo and Taipale, 2008), but *Indian hedgehog (IHH)* has been shown to be highly expressed in human colon (Van den Brink, 2007; van den Brink et al., 2004). The presence and role, if any, of *Desert hedgehog (DHH)* has not been characterized in the colon, although *DHH* has been linked to maintenance and regeneration of the corneal epithelium (Kucerova et al., 2012).

Thus far, only three studies have detailed the importance of epithelial Wnts in homeostasis or response to injury in the intestine (O’Connell et al., 2018; Suh et al., 2017; Zou et al., 2018), with most studies focused on the role of mesenchymal Wnts in homeostasis and disease (Gregorieff et al., 2005; Greicius et al., 2018; Koch, 2017; Shoshkes-Carmel et al., 2018; Valenta et al., 2016). The majority of data gained on mouse intestinal injury models suggests that the mesenchymal Wnts are necessary for epithelial regeneration, but did not characterize the role epithelial Wnts may be playing in these processes.

Human colonoid cultures are a tractable, epithelial-only model that can indefinitely proliferate due to the presence of adult intestinal stem cells (Sato et al., 2011), making them an excellent model to study intestinal crypt injury and hyperplasia. Foodborne bacterial pathogens, such as enterohemorrhagic *E. coli* (EHEC) or *Citrobacter rodentium*, a mouse-adapted bacterium that affects the intestine similarly to EHEC, can cause severe damage to the intestinal epithelia, resulting in hyperproliferation and crypt hyperplasia post-infection (Khan et al., 2006; Vallance et al., 2003; Xicohtencatl-Cortes et al., 2007). We have previously characterized the EHEC-secreted serine protease cytotoxin, EspP, as an important virulence factor in EHEC infection and colonic epithelial damage (In et al., 2013). Cytotoxins in the family of <u>s</u>erine <u>p</u>rotease <u>a</u>uto<u>t</u>ransporters of *<u>E</u>nterobacteriaceae* (SPATEs) are secreted by most pathogenic *E. coli* and have well characterized functions that aid in bacterial adherence and colonization of epithelial cells (Dautin, 2010). Two SPATEs, Pet and EspC, secreted by enteroaggregative *E. coli* and enteropathogenic *E. coli*, respectively, cause cytotoxicity to intestinal explants (Henderson et al., 1999; Mellies et al., 2001). However, whether or not EspP has cytotoxic properties on intestinal cells has been controversial (Weiss and Brockmeyer, 2012).

In this study, we used the EHEC cytotoxin, EspP to induce epithelial injury and model the intestinal stem cell response that includes the initiation of regeneration using stem cell-derived human colonoids. Using both molecular and proteomics-based approaches, we found that epithelial-produced WNT2B and Desert Hedgehog-activated Hedgehog signaling interact and are necessary for human colonoid regeneration.

## RESULTS

### EspP, a bacterial autotransporter, has a serine protease-dependent cytotoxic effect on human colonoids

To determine if EspP induces cytotoxicity in a serine protease-dependent manner in human colonoids, we added recombinant EspP or its serine protease-deficient mutant, EspP 263A (Khan et al., 2011), to normal human colonoids. After an overnight treatment with EspP (50 μg/ml), all human colonoid lines used in this study (Supplementary Table 1) exhibited cell shedding and loss of colonoid structure, indicators of cell death (Figure 1). In contrast, overnight treatment with the protease-deficient mutant, EspP S263A (50 μg/ml), had no visible detrimental effect on the colonoids. Therefore, EspP has a cytotoxic effect on human colonoids and this activity is serine protease-dependent.

**Table 1.** EspP-injured colonoids upregulated proteins in the Wnt and Hedgehog pathways. Selected proteins from the proteomics analysis show that proteins in the Wnt and Hedgehog pathways are upregulated in the EspP-injured colonoids. The ratio is protein expression of EspP-injured over control colonoids.

| **Accession** | **Description ( Homo sapiens)** | **Coverage** | **Ratio**<br>**(EspP/control)** |
| --- | --- | --- | --- |
| **630044901** | **Protein Wnt2b isoform 3** | **4.35** | **3.101** |
| **27883842** | **Polycomb complex protein BMI-1** | **2.15** | **2.057** |
| **4502805** | **chromogranin-A isoform 1 preproprotein** | **39.61** | **2.009** |
| **31542745** | **protein wntless homolog isoform 1 precursor** | **6.10** | **1.997** |
| **24431935** | **reticulon-4 isoform A** | **12.08** | **1.995** |
| **4506055** | **cAMP-dependent protein kinase catalytic subunit**<br>**alpha isoform 1** | **36.47** | **1.807** |
| **20544145** | **casein kinase I isoform delta isoform 2** | **16.63** | **1.734** |
| **395394053** | **disheveled-associated activator of morphogenesis 1**<br>**isoform 2** | **7.49** | **1.585** |
| **225903437** | **glycogen synthase kinase-3 beta isoform 2** | **20.95** | **1.678** |
| **395394053** | **disheveled-associated activator of morphogenesis 1**<br>**isoform 2** | **7.49** | **1.585** |
| **188528675** | **slit homolog 1 protein precursor** | **4.69** | **1.520** |
| **34485714** | **ras-related protein Rab-23** | **12.24** | **1.477** |
| **33636738** | **cAMP-dependent protein kinase catalytic subunit**<br>**beta isoform 1** | **28.14** | **1.464** |
| **25121993** | **RNA-binding protein Musashi homolog 2 isoform b** | **27.49** | **1.391** |
| **148727288** | **low-density lipoprotein receptor-related protein 6**<br>**precursor** | **4.15** | **1.350** |
| **14916475** | **protein Wnt-3a precursor** | **27.56** | **1.269** |
| **578808446** | **PREDICTED: slit homolog 2 protein isoform X5** | **13.65** | **1.259** |
| **4885523** | **noggin precursor** | **22.84** | **1.247** |
| **578808417** | **PREDICTED: prominin-1 isoform X5** | **3.68** | **1.194** |
| **339276103** | **R-spondin-1 isoform 3 precursor** | **31.50** | **1.167** |
| **11545873** | **SPARC-related modular calcium-binding protein 1**<br>**isoform 2 precursor** | **7.14** | **1.141** |

**Figure 1.**
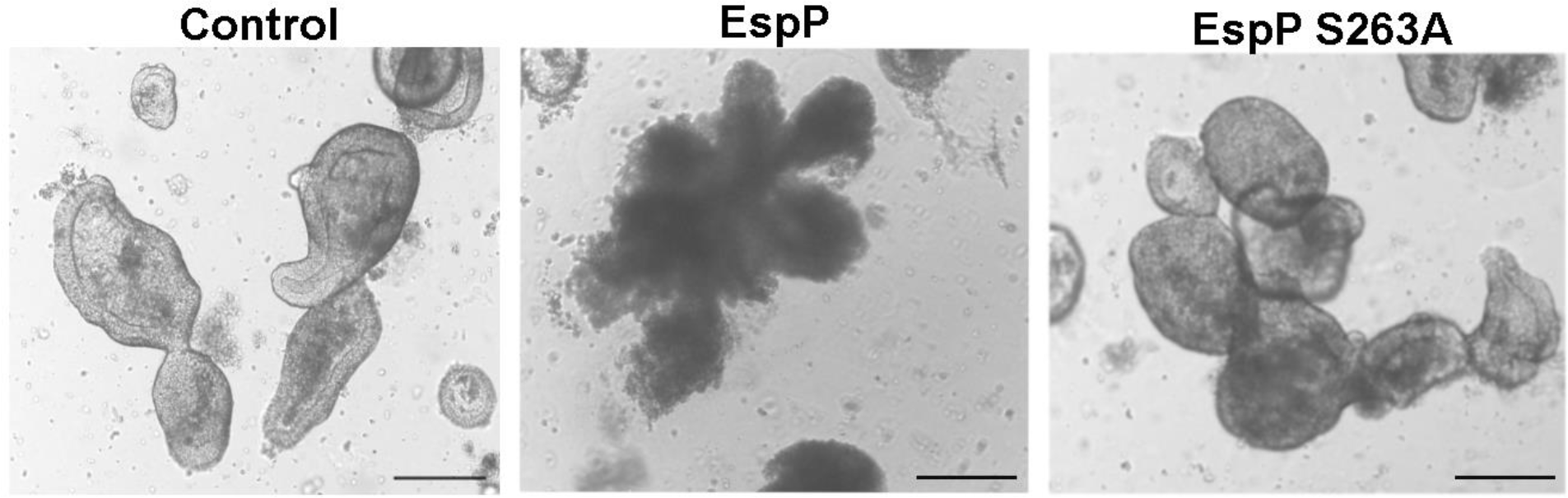
EspP requires serine protease function to cause cytotoxicity of human colonoids. Representative images of colonoids after overnight treatment: control (left), EspP-treated (middle), and EspP 263A-treated (right). EspP requires serine protease activity to have a cytotoxic effect on the colonoids; scale bar = 200 μm. N=3

We hypothesized that EspP-induced injury would model the EHEC-induced denuded colonic epithelia and crypt hyperplasia, the latter mimicked by colonoid regeneration after EspP-induced injury. To test this hypothesis, control and EspP-injured colonoids were harvested after overnight EspP treatment and replated to monitor for colonoid regeneration. The formation of colonoids in the EspP-treated cultures was observed at 24h and 48h post-replating (Figure 2A). At 24h, the colonoids were generally smaller in size compared to control and primarily spheroids. In contrast, at 48h, the regenerating colonoids more resembled the control culture, with colonoids beginning to form multi-lobular structures (Figure 2B). Therefore, human colonoids can regenerate after injury by the bacterial cytotoxin EspP.

**Figure 2.**
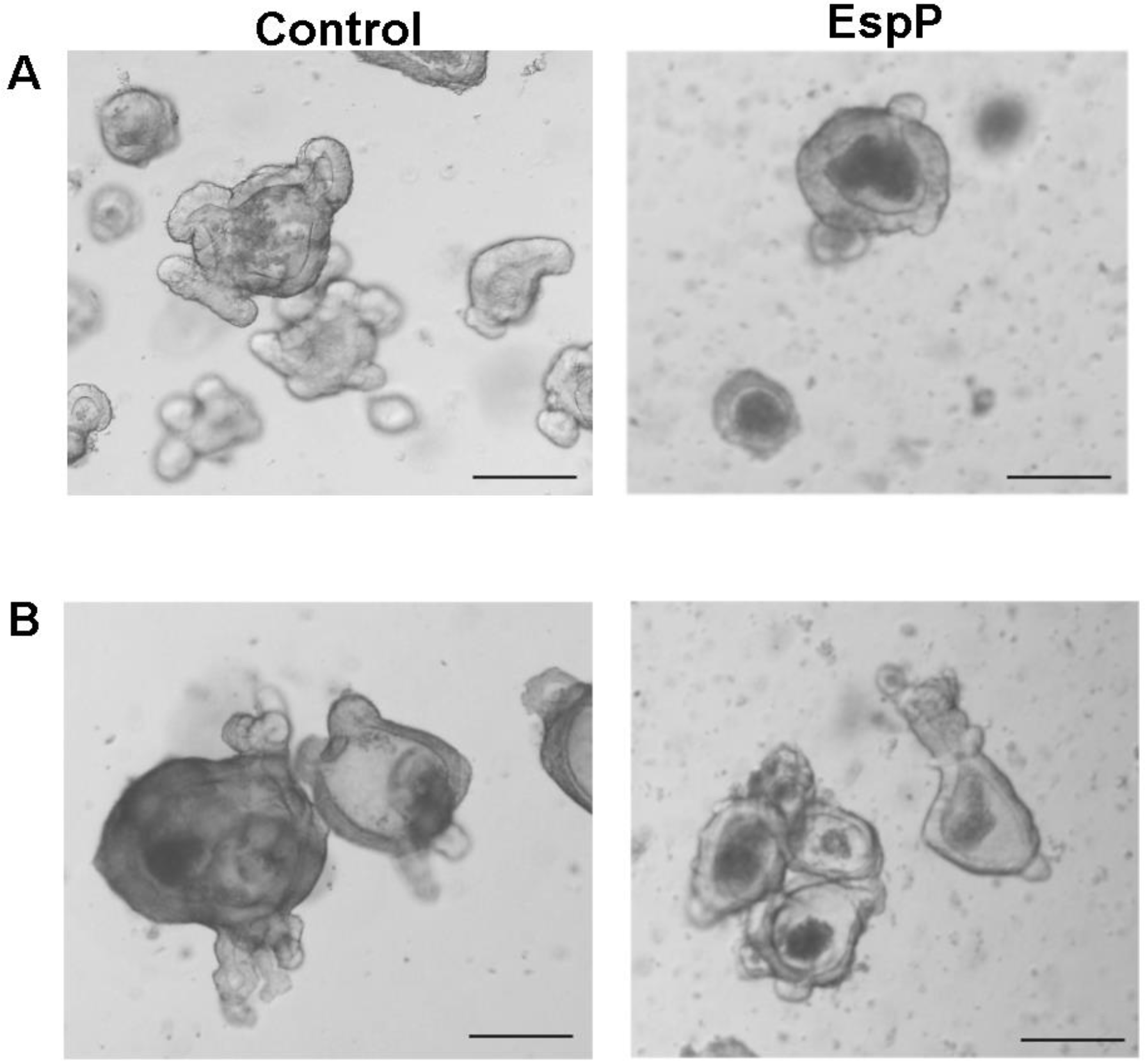
Colonoids can model crypt regeneration after EspP washout. (A and B) Representative images of colonoids after washout and replating: (A) 24h post-washout and (B) 48h post-washout; scale bar = 200 μm. N>3

### Proteomics analysis shows WNT2B and Desert Hedgehog are upregulated during regeneration

To begin to identify key regeneration-associated pathways, we employed a proteomics approach. Control, EspP-and EspP S263A-treated colonoids were harvested, lysed, and the proteins were identified and quantified with tandem mass spectrometry and iTRAQ. Over 5000 proteins in the EspP-treated culture were found up- or down-regulated compared to the control culture, with very little overlap of differentially expressed proteins between the EspP- and EspP S263A-treated cultures (Figure S1A). The majority of proteins identified in the proteomics assay were cytosolic or nuclear (Figure S1B). A key group of proteins that was upregulated in the EspP treated cultures were those associated with Wnt, Hedgehog, and putative stem cell-related proteins. An abbreviated list of these proteins is shown in Table 1. WNT2B isoform 3, WNT3A, Wntless and numerous downstream targets of GLI1 (active hedgehog signaling) were upregulated in the EspP-injured culture. Many of the proteins listed in Table 1 were either not changed or downregulated in the EspP S263A-treated (no cytotoxicity) culture suggesting that EspP specifically induced activation of Wnt and Hedgehog signaling as part of the colonic damage and regenerative response.

To assess WNT2B expression in the colonoids post-EspP injury, we performed immunostaining. WNT2B expression was concentrated in specific, rare epithelial cells in normal human colon crypt (Figure S2A) and in colonoids (Figure S2B). Not every colonic crypt or every colonoid had WNT2B-positive cells. However, colonoids regenerating 24h after EspP-induced injury contained more WNT2B+ cells and diffuse WNT2B staining throughout the colonoid (Figure S2C and C’).

We performed qRT-PCR to validate the key pathway molecules identified in the proteomics screen. The mRNA expression of select stem cell, Wnt, and Hedgehog genes was compared between EspP-injured regenerating (at the 24h timepoint) and control colonoids. Although the injured colonoids regenerate to re-form their 3D structure after EspP washout, the intestinal stem cell markers *LGR4* and *LGR5* were not upregulated. *LGR4* was significantly downregulated, whereas *LGR5* was unchanged (Figure 3). *BMI1*, which was significantly upregulated in the proteomics result, showed an upward trend in its mRNA expression, but without reaching statistical significance. The proteomics screen identified WNT2B isoform 3 as significantly upregulated in the EspP-injured regenerating colonoids. The EspP-injured regenerating colonoids had a slight downregulation of *WNT2B2* (previously known as *WNT13A*), an upward trend of *WNT2B1* (*WNT13B*), and a significant upregulation of *WNT2B3* (*WNT13C*) (Figure 3). WNT2B3’s upregulation following EspP-induced injury confirmed the proteomics assay, but was still unexpected as it is not thought to be a classical epithelial-produced Wnt.

**Figure 3.**
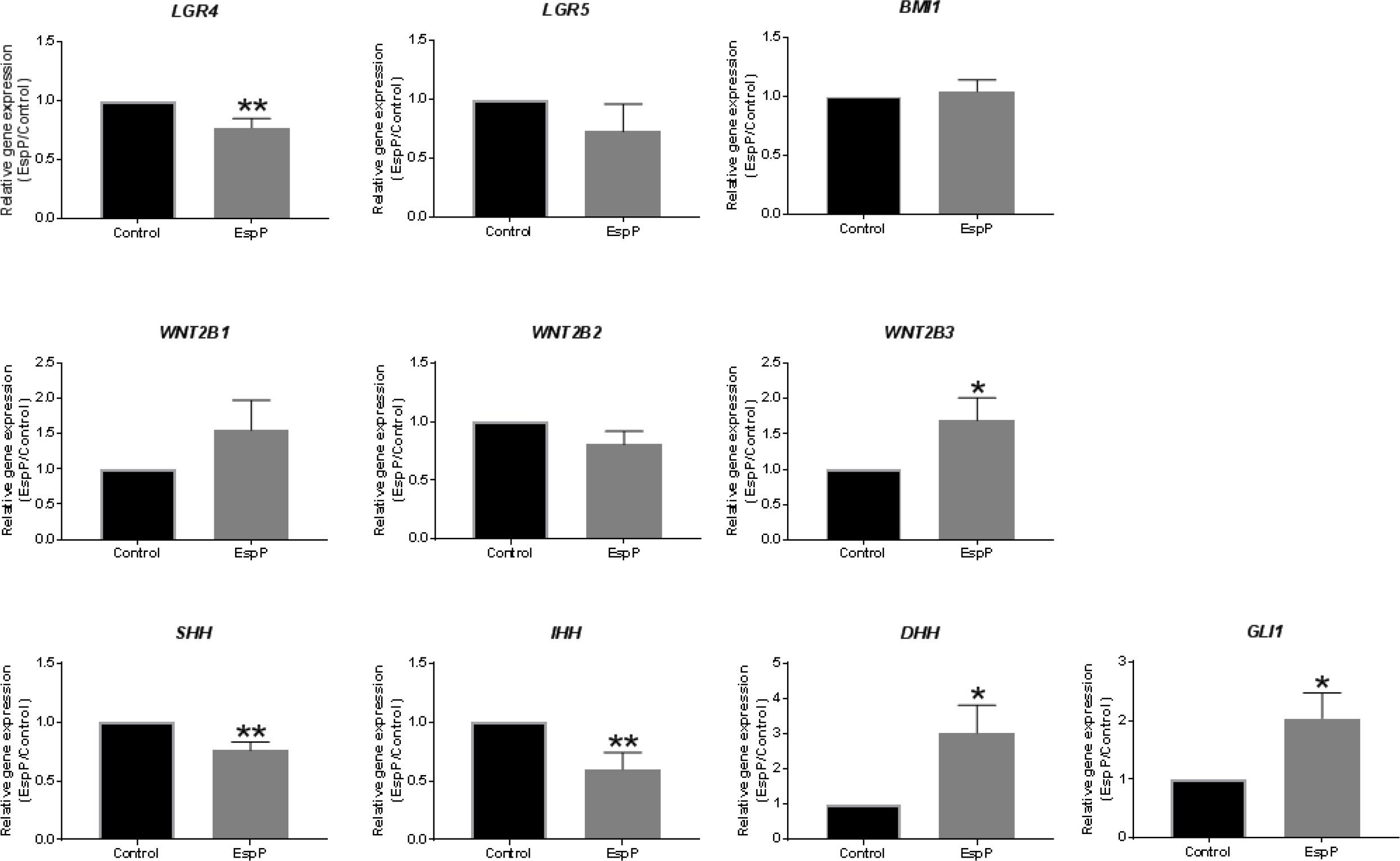
EspP-treated colonoids upregulate WNT2B and DHH during regeneration. Gene expression of regenerating colonoids was analyzed by qRT-PCR. Relative gene expression is shown as a ratio of EspP-treated compared to control colonoids, and normalized to 18S. * p < 0.05; ** p < 0.01. N≥11

Since numerous downstream targets of Hedgehog signaling were upregulated in the regenerating colonoids, we evaluated whether the canonical Hedgehog effectors *GLI1* and *GLI2* were changed in the regenerating colonoids. Both genes have been found upregulated in colon cancer cell lines (Mazumdar et al., 2011; Zhang et al., 2017) and implicated in cancer cell proliferation. *GLI2* transcripts were not detected in either the control or regenerating colonoids. However, *GLI1* was significantly upregulated in the regenerating colonoids (Figure 3). Only the hedgehog ligand *DHH* was significantly upregulated in the regenerating colonoids (Figure 3). Both hedgehog ligands *SHH* and *IHH* were significantly downregulated. Overall, the EspP-injured regenerating colonoids lead to upregulation of hedgehog signaling, specifically via the hedgehog ligand, DHH.

### Epithelial Wnt is indispensable for colonoid regeneration

To determine whether epithelial produced Wnts are important for colonoid regeneration, control and EspP-injured colonoids were monitored in the absence (Figure 4A) or presence (Figure 4B) of IWP-2 (2.5 μM), a porcupine inhibitor that inhibits palmitoylation of all Wnts and results in inhibition of processing and secretion of Wnts (Farin et al., 2012). The colonoid media containing 50% v/v Wnt3A conditioned media was maintained in all experimental conditions. As previously shown, the EspP-injured colonoids were able to regenerate and re-form 3D colonoids after EspP is removed (Figure 4A). In the continued presence of IWP-2 (pre-treatment prior to EspP addition, during EspP treatment, and during the 24h regeneration period), EspP-injured colonoids were unable to re-form 3D colonoids. Interestingly, the control culture showed no morphologic difference in the presence of IWP-2 (Figure 4B). This suggests that the Wnt3A conditioned media is sufficient to maintain homeostatic growth and proliferation of colonoids, but is not sufficient for regeneration following EspP-induced injury. Inhibition of epithelial Wnt secretion (by IWP-2) prevents human colonoid regeneration. This indicates that epithelial Wnt(s) are necessary for regeneration.

**Figure 4.**
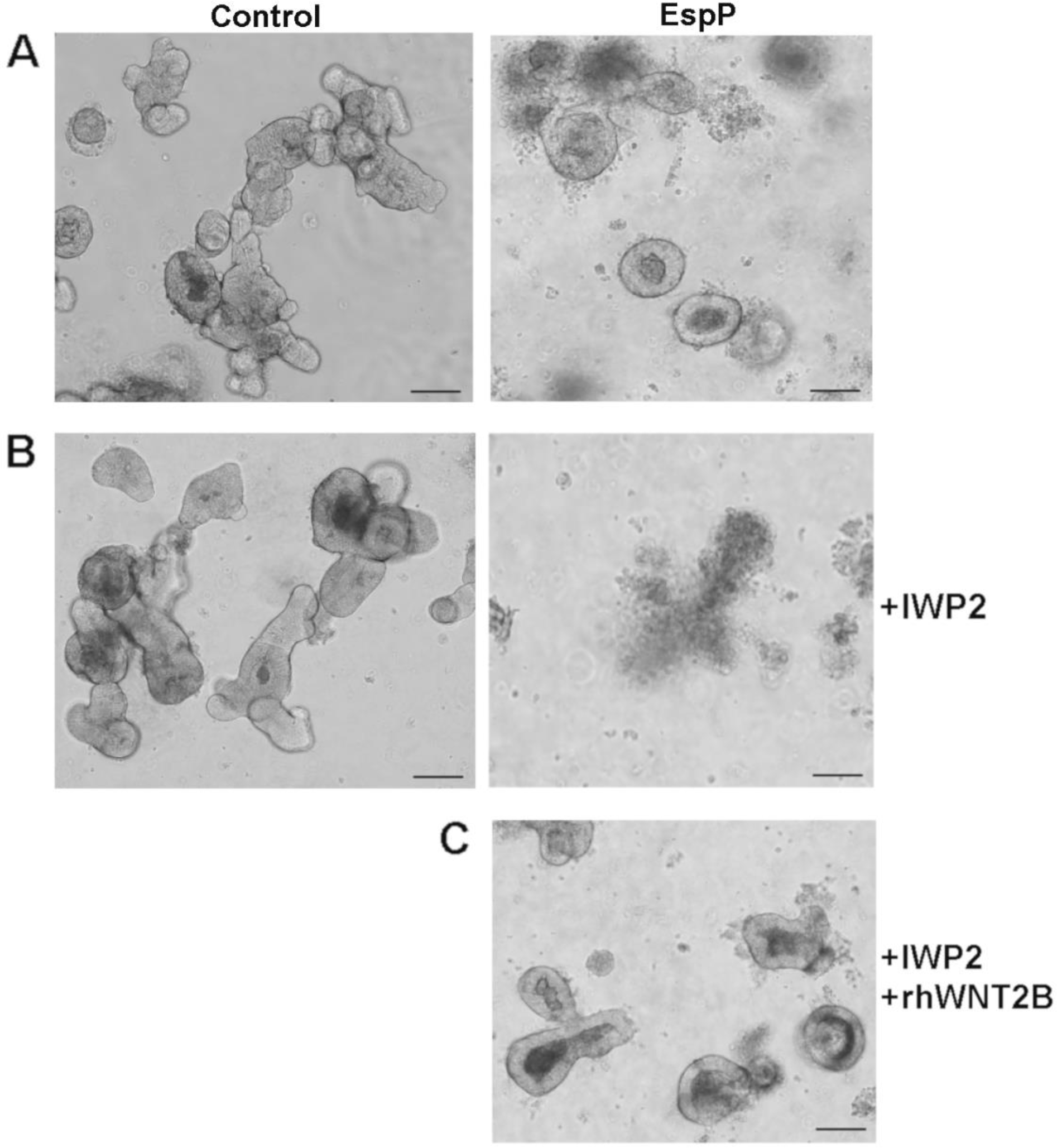
WNT2B compensates for inhibition of epithelial wnts in regenerating colonoids. (A - C) Control (left) and EspP-treated (right) colonoids after washout, at 24h regeneration. (A) Representative images of colonoids at 24h regeneration. (B) Representative images of colonoids in the continued presence of IWP2 at 24h regeneration. Representative image of colonoids in the continued presence of IWP2 and recombinant human WNT2B (rhWNT2B) at 24h regeneration; scale bar = 200 μm. N≥4

The proteomics screen identified upregulation of WNT2B3 in the EspP-injured colonoids. We evaluated if WNT2B alone could stimulate regeneration. Recombinant human WNT2B (rhWNT2B) was added to colonoids at the same time as IWP-2 and kept in the cultures during the course of the experiment. Although IWP-2 inhibited colonoid regeneration, rhWNT2B was sufficient to rescue and promote regeneration after EspP-induced injury (Figure 4C). To determine the direct effect of epithelial WNT2B on colonoid regeneration post EspP-injury, we used a lentiviral shRNA approach to knockdown *WNT2B* in the colonoids. As a technical control, we used a lentiviral shRNA to knockdown *DRA (SLC26A3)* in human duodenal enteroids. At 17 days post-transduction (15 days after the start of puromycin selection), the enteroids with *DRA* shRNA were thriving in the presence of puromycin (Figure S3). In contrast, the colonoids with *WNT2B* shRNA sharply declined and were unable to propagate (Figure S3). This result is consistent with the report by O’Connell et al., 2018 in which the enteroids and colonoids derived from WNT2B-deficient individuals were not stable and could only form a short-term culture in the presence of recombinant murine WNT2B.

Studies in chick retinal explants found that Wnt2b overexpression leads to increased cell proliferation and the growth of large, folded retinal tissue (Ohta et al., 2011). However, co-overexpression of Wnt2b with the small, leucine-rich proteoglycan Tsukushi (Tsk) led to an inhibition of the Wnt2b-dependent hyperproliferation. Since we could not create a viable *WNT2B* KD human colonoid line, we examined whether TSK could inhibit WNT2B function in colonoids. Colonoids were treated with recombinant human TSK (rhTSK). Similar to the presence of IWP-2, control colonoids showed no morphologic difference in the presence of rhTSK (Figure 5). However, the EspP-injured colonoids were unable to regenerate in the presence of rhTSK. Taken together, these data indicate that epithelial WNT2B is necessary for colonoid regeneration after EspP-induced injury.

**Figure 5.**
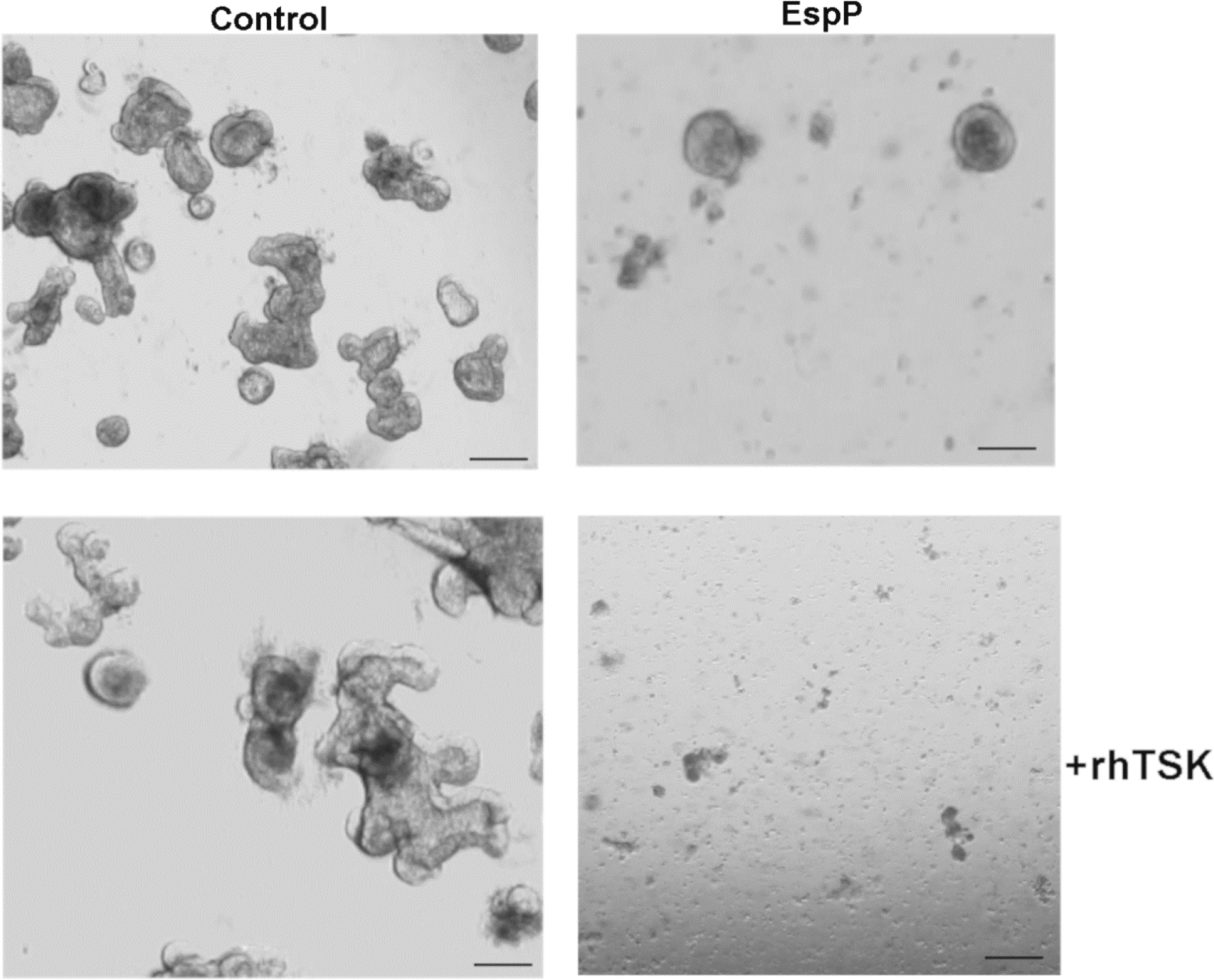
TSK inhibits colonoid regeneration post EspP-treatment. Representative images of control (left) and EspP-treated (right) colonoids after washout, at 24h regeneration. Control and EspP-treated colonoids were in the continued presence of recombinant human Tsukushi (rhTSK) (bottom panel); scale bar = 200 μm. N=3

### DHH activated hedgehog signaling modulates WNT2B

The regenerating colonoids also had a significant upregulation of *DHH* and *GLI1* (Figure 3) suggesting an active role for hedgehog signaling following EspP-induced injury. To determine whether there was a link between Hedgehog signaling and WNT2B in regeneration, we treated colonoids with either the Smoothened agonist (SAG) or recombinant human DHH (rhDHH prior to EspP exposure. SAG binds to Smoothened and induces activation of the Hedgehog pathway (Chen et al., 2002). Its function is thought to be Hedgehog ligand-independent. DHH, as a Hedgehog ligand, also activates the Hedgehog pathway. Colonoids present 24h after regeneration were collected and analyzed for gene expression of stem cell markers, WNT, and Hedghog pathway molecules. mRNA expression in the presence of the agonists was compared to control (no agonists). The intestinal stem cell markers *LGR4* and *LGR5* were further downregulated in the presence of SAG compared to control. However, both genes were upregulated in the presence of rhDHH compared to control (Figure 6). *BMI1* remained largely unchanged with SAG treatment, but was significantly upregulated in the EspP-injured, rhDHH treated colonoids, similar to the upregulation of *LGR4* and *LGR5*. This suggests that DHH activates a specific Hedgehog pathway that SAG does not. DHH-activated signaling has a direct effect on the intestinal stem cell markers.

**Figure 6.**
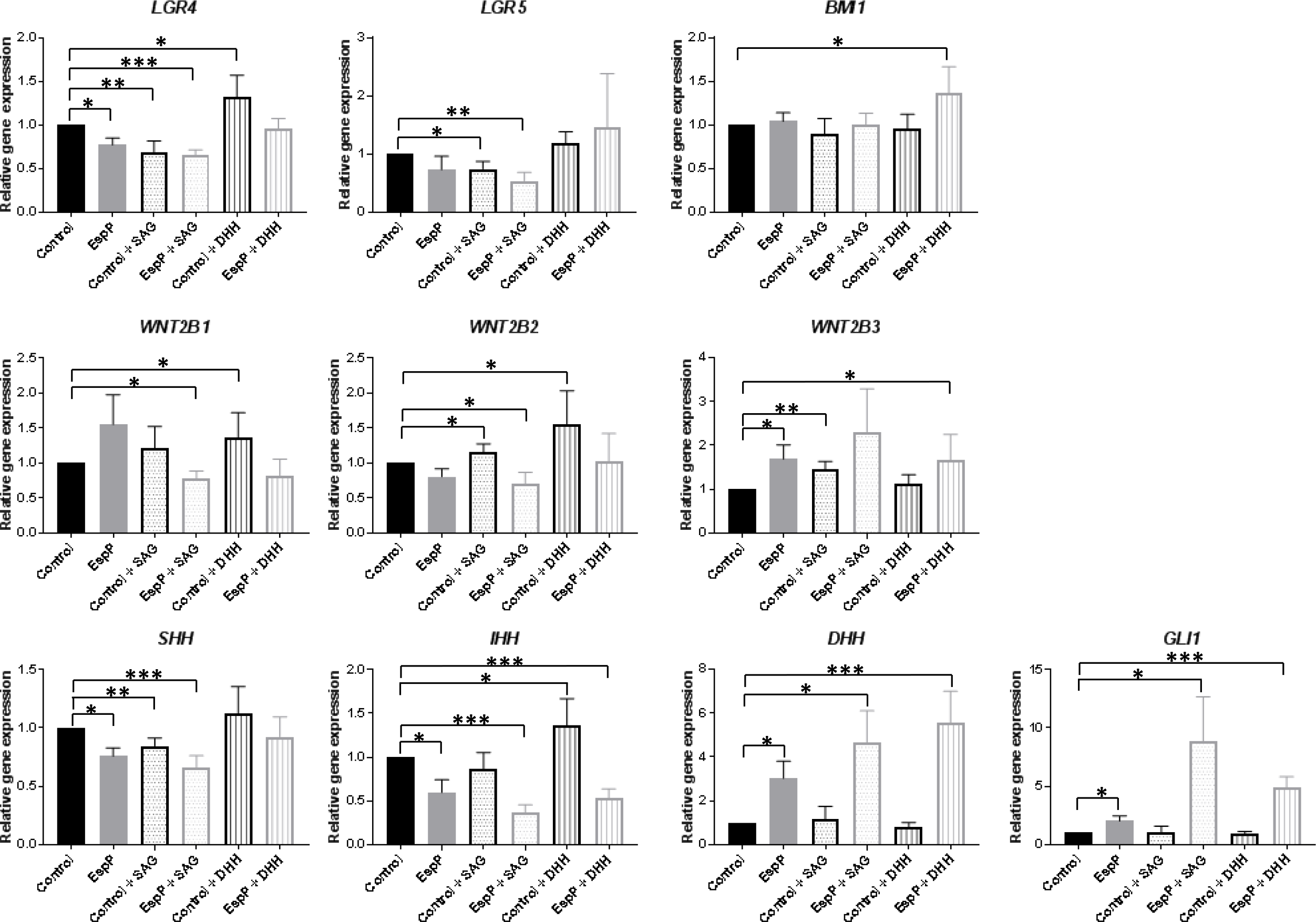
Hedgehog agonists upregulate WNT2B3 and DHH during regeneration. Control and EspP-treated colonoids were treated with Smoothened agonist (SAG) or recombinant human Desert Hedgehog (DHH). Gene expression of regenerating colonoids was analyzed by qRT-PCR. Relative gene expression is shown as a ratio of treated (EspP and/or SAG or DHH) compared to control colonoids, and normalized to 18S. n>3; * p < 0.05; ** p < 0.01; *** p < 0.001. N≥3

SAG treatment significantly downregulated *WNT2B1* and *WNT2B2* in EspP-injured colonoids. In contrast, *WNT2B3* expression continued to trend upwards. rhDHH treatment had no effect on *WNT2B1* and *WNT2B2* expression in EspP-injured colonoids. However, *WNT2B3* was significantly upregulated (Figure 6). This suggests that DHH positively modulates WNT2B3 expression during colonoid regeneration. Similar to the stem cell markers that we evaluated, SAG and rhDHH caused different expression patterns of the three *WNT2B* isoforms.

SAG treatment either significantly downregulated or had no effect on expression of *SHH* and *IHH* in both control and EspP-injured colonoids, but significantly upregulated *DHH* and *GLI1* expression in EspP-injured colonoids. EspP-injured colonoids showed significant upregulation of *GLI1* and *DHH* in the presence of rhDHH, compared to control (Figure 6). These data show that both SAG (hedgehog activation) and rhDHH can modulate *WNT2B* expression, but only *WNT2B3* is upregulated in EspP-injured colonoids with these hedgehog agonists. DHH acts in a specific manner to activate hedgehog signaling following injury to the colonoids. Although SAG and rhDHH treatment similarly upregulated *DHH* and *GLI1* in EspP-injured colonoids, they had different effects on the expression levels of the other genes interrogated. This suggests that DHH activates Hedgehog signaling in a SAG-independent manner. Overall, these results show that human colonoids regenerate after bacterial cytotoxin-induced injury via interaction of the DHH and WNT2B-dependent pathways.

## DISCUSSION

Intestinal regeneration is dependent on Wnt signaling to stimulate stem cell proliferation. Most studies have focused on the identity of the intestinal stem cells that drive proliferation and crypt hyperplasia in mouse models under both normal and post-injury conditions, particularly post-radiation (Hua et al., 2012; Hua et al., 2017; Kuruvilla et al., 2016; Metcalfe et al., 2014; Zhou et al., 2013). The regenerative pathways and key players in these pathways are not well understood. In this study, we focused on characterizing the molecules that drive a regenerative response following exposure to a virulence factor in a bacterial diarrheal disease: EspP, an EHEC-secreted bacterial cytotoxin that causes epithelial damage. Colonic regeneration is dependent on epithelial signals, namely WNT2B and DHH. These two molecules activate Wnt and Hedgehog signaling interaction during colonic regeneration.

Using the human colonoid model, which contains no mesenchyme, we employed a proteomics screen to characterize the pathways that are active following EspP-induced injury. WNT2B and numerous proteins downstream of active Hedgehog signaling were upregulated, suggesting Wnt and Hedgehog signaling are important in colonoid regeneration. Both pathways have been implicated in organ development and maintenance (Clevers, 2006; Petrova and Joyner, 2014), with Hedgehog signaling described as important in regeneration of most organs (Adolphe, 2004; Konstantinou et al., 2016; Langiewicz et al., 2016; Le et al., 2008; Sriperumbudur et al., 2016; Trowbridge et al., 2006; Wang et al., 2016a; Wang et al., 2016b). Although recent studies have focused on the crosstalk between Wnt and Hedgehog signaling in cancer progression (Jiang et al., 2014; Regan et al., 2017; Song et al., 2015), these two pathways also have been implicated in regeneration of bladder epithelia, bone, and adrenal glands (Day and Yang, 2008; Finco et al., 2018; Shin et al., 2011).

Our results indicate that epithelia-produced WNT2B and DHH are important regulators of human colonoid regeneration, with DHH modulating *WNT2B3* expression following EspP-induced injury. Activation of this particular Hedgehog pathway is not redundant between the three mammalian Hedgehog ligands. Sonic and Indian Hedgehog transcripts were either downregulated or unchanged during regeneration. Most of our understanding of Hedgehog signaling focuses on Sonic Hedgehog, likely because it is the most widely expressed mammalian Hedgehog ligand (Varjosalo and Taipale, 2008). The implications of downregulated *SHH* in colonoid regeneration are not clear, however, previous studies have shown that *IHH* downregulation initiates intestinal wound healing and abrogates adenoma development (Büller et al., 2015; van Dop et al., 2010). Until now, DHH function has not been well understood. It is primarily described as an essential factor in gonad (O’Hara et al., 2011; Rothacker et al., 2018; Yao et al., 2002) or peripheral nerve development (Bajestan et al., 2006; Parmantier et al., 1999). However, one study demonstrated an essential role for DHH in corneal homeostasis and regeneration (Kucerova et al., 2012). Our results highlight a novel role for DHH-activated Hedgehog signaling in human colonic regeneration.

In human colonoids and colonic tissue, WNT2B is localized to a rare cell that is not present in every colonoid or crypt. The identity of this cell in human colonoids is currently unknown but under further investigation. Regeneration following cytotoxin-induced injury results in diffuse WNT2B staining with a higher number of WNT2B+ cells, similar to a study that showed upregulation of Wnt2b in mouse intestinal crypts post-irradiation (Suh et al., 2017). This correlates with the upregulation of *WNT2B3* mRNA in the regenerating colonoids. Although WNT2B has been characterized as having two isoforms in cancer cells (Katoh, 2001), three WNT2B isoforms have been identified in multiple mammalian cells and been shown to function disparately from each other (Bunaciu et al., 2008). Since our proteomics screen identified the WNT2B isoform 3, we used the primers described by Bunaciu et al. to distinguish between the WNT2B isoforms. The three isoforms were regulated differently during regeneration and in the presence of Hedgehog agonists, SAG and rhDHH.

Mesenchymal Wnts are clearly essential for regeneration (Gregorieff et al., 2005; Greicius et al., 2018; Koch, 2017; Shoshkes-Carmel et al., 2018; Valenta et al., 2016), but only a few studies have highlighted the importance of epithelial Wnts in intestinal development and injury response (O’Connell et al., 2018; Suh et al., 2017; Zou et al., 2018). Of note, recently *WNT2B* mutations were found to cause neonatal-onset chronic diarrhea, with inflammation seen in the stomach, duodenum, and colon (O’Connell et al., 2018). This study showed that enteroids from these *WNT2B*-deficient patients could not form stable cultures, although addition of recombinant murine Wnt2b stabilized the cultures for a short period. This study emphasizes the significant differences between the regeneration potential of mouse and human intestinal epithelium. Knockout or knockdown of *Wnt2b* in the whole mouse or mouse organoids, respectively, results in no detrimental phenotype. However, human intestinal epithelial WNT2B is indispensable in intestinal development and regeneration following injury. Taken together, our studies indicate that data gained on mouse models of intestinal development, homeostasis, and injury may not directly translate to human intestinal physiology and pathophysiology.

In summary, using the bacterial cytotoxin EspP to model damage, we showed that human colonoids can be used to study the role of epithelial molecules in regeneration. Epithelial WNT2B and Desert Hedgehog are essential and interact during regeneration following injury. Importantly, the hedgehog ligands, Desert, Indian, and Sonic, are not redundant in colonic regeneration. Understanding the mechanisms that specifically drive WNT2B3 and DHH in colonic development and regeneration may provide the basis for useful therapeutics in controlled regeneration in patients with some colonic diseases.

## EXPERIMENTAL PROCEDURES

### Tissue collection and colonoid generation

Colonic biopsies from healthy individuals were obtained under Johns Hopkins University School of Medicine Institutional Review Board (IRB#NA_00038329) and are detailed in Supplementary Table 1. Colonic crypt isolation and colonoid generation were prepared as previously reported (In et al., 2016; Jung et al., 2011). Briefly, biopsy tissue was minced, washed several times in freshly prepared cold chelating solution (CCS; 5.6mM Na2HPO4, 8mM KH2PO4, 96.2mM NaCl, 1.6mM KCl, 43.4mM sucrose, 54.9mM D-sorbitol, and 0.5mM DL-dithiothreitol) and incubated 1 hour at 4°C in 10 mM EDTA in CCS on an orbital shaker. Isolated crypts were resuspended in Matrigel (Corning, Tewksbury, MA) and 30 ul droplets were plated in a 24-well plate (Corning). After polymerization at 37°C, 500 ul of expansion media (EM) was added for 2 days (Advanced Dulbecco’s modified Eagle medium/Ham’s F-12 (ThermoFisher, Waltham, MA), 100 U/mL penicillin/streptomycin (Quality Biological, Gaithersburg, MD), 10 mM HEPES (ThermoFisher), and 1X GlutaMAX (ThermoFisher), with 50% v/v WNT3A conditioned medium (ATCC CRL-2647), 15% v/v R-spondin1 conditioned medium (cell line kindly provided by Calvin Kuo, Stanford University), 10% v/v Noggin conditioned medium (cell line kindly provided by Gijs van den Brink, Tytgat Institute for Liver and Intestinal Research), 1X B27 supplement (ThermoFisher), 1mM N-acetylcysteine (MilliporeSigma), 50 ng/mL human epidermal growth factor (ThermoFisher), 10 nM [Leu-15] gastrin (AnaSpec, Fremont, CA), 500 nM A83-01 (Tocris, Bristol, United Kingdom), 10 μM SB202190 (MilliporeSigma), 100 mg/mL primocin (InvivoGen, San Diego, CA), 10 μM CHIR99021 (Tocris), and 10 μM Y-27632 (Tocris)). After 2 days, the EM (without CHIR99021 and Y-27632) was replaced every other day. Colonoids were passaged every 7 days by harvesting in Cultrex Organoid Harvesting Solution (Trevigen, Gaithersburg, MD) at 4°C with shaking for 30.’ Colonoids were fragmented by trituration with a P200 pipet 30-50 times, collected and diluted in Advanced DMEM/F12, centrifuged at 300 xg for 10’ at 4°C. The pellet was resuspended in Matrigel and plated as described for crypt isolation. All colonoid cultures were maintained at 37°C and 5% CO_2_. Unless noted, colonoid lines have been passaged >20 times.

### Recombinant EspP generation and collection

AD202 cells transformed with the plasmid encoding wild-type EspP (pRLS5) and serine protease-deficient mutant EspP S263A was kindly provided by H. Bernstein, NIH (Szabady et al., 2004). The cells were grown at 37°C in Luria-Bertani (LB) broth (ThermoFisher), overnight. They were then pelleted, washed, and grown at 37°C in fresh LB broth for approximately 15’. IPTG (100 μM) was added to induce *espP* or *espP S263A* expression. The culture was grown until reaching an OD_550_ 2.0. Bacterial cells were removed by centrifugation (9000 rpm, 30’, 4°C, Sorvall RC6, SLA-3000 rotor). EspP and EspP S263A was collected from the cell-free supernatant by ammonium sulfate precipitation (60%, o/n, 4°C), followed by centrifugation (9000 rpm, 30’, 4°C, Sorvall RC6, SLA-3000 rotor). The pellet was resuspended in PBS, syringe filtered (0.2 μm), then diluted with 15% glycerol to allow for freezing. Each batch of recombinant EspP and EspP S263A was separated on SDS-PAGE and stained with Coomassie Blue to check purity. Protein concentrations were determined by Bradford assay (Bio-Rad, Hercules, CA). Serine protease activity was determined by pepsin A-cleavage assay (Brockmeyer et al., 2007).

### EspP treatment and colonoid regeneration

Colonoids were plated in Matrigel in 24 well plates and separated into experimental conditions (control, EspP treatment, EspP plus inhibitors or agonists). Since the mechanics of passaging colonoids includes fragmenting their 3D structure and therefore causing injury, we attempted to minimize this by not triturating the colonoids, but instead, harvesting them without fragmentation and replating into new Matrigel. Colonoids were pre-treated with inhibitors or agonists at least 8h prior to overnight EspP treatment. After overnight treatment, colonoids were harvested in Cultrex Organoid Harvesting Solution, washed twice in Advanced DMEM/F12, and pelleted at 300 xg for 10’ at 4°C, and replated in Matrigel for 24h regeneration. After replating, colonoids were kept in the presence of any inhibitors or agonists using during the experiment. After the 24h regeneration period, colonoids were imaged or processed for further studies. All experimental reagents used are detailed in Supplementary Table 2.

### Brightfield imaging

Colonoids plated in Matrigel in 24 well plates were imaged during the course of experiments on a Zeiss Axio Observer A1 inverted microscope (Zeiss, Oberkochen, Germany) with images captured on CellSense imaging software (Olympus, Tokyo, Japan). Images were viewed and processed using OlyVia (Olympus).

### Immunofluorescence staining and confocal imaging

Fixed tissues were frozen in OCT and sectioned (10 μm thick). Colonoids were harvested from Matrigel using Cultrex Organoid Harvesting Solution. They were pelleted (300 xg, 10’, 4°C), and fixed for 40 min in 4% paraformaldehyde (Electron Microscopy Sciences, Hatfield, PA). Both fixed tissue and colonoids were permeabilized and blocked simultaneously for 1h using a 10% Fetal Bovine Serum (Atlanta Biologicals, Flowery Branch, GA), 0.1% saponin (MilliporeSigma) solution prepared in PBS. After three PBS washes, 100 μl of primary antibody against WNT2B (HPA060696, MilliporeSigma) prepared at 1:100 dilution in PBS was added to the cells and incubated overnight at 4°C. Afterwards, cells were washed 3 times with PBS, and 100 μl of AlexaFluor secondary antibodies, AlexaFluor-647 phalloidin, and Hoechst 33342 (1 mg/ml, all ThermoFisher), diluted 1:100 in PBS, were added for 1h at room temperature. After three PBS washes, 50 μl of FluorSave Reagent (Calbiochem) was added to the colonoids and they were mounted between a glass slide and a number 1 coverslip. Confocal imaging was carried out in the Imaging Core of the Hopkins NIH/NIDDK Basic and Translational Research Digestive Disease Core Center using a LSM510 META laser scanning confocal microscope running ZEN 2012 (black edition) imaging software (Zeiss).

### Protein extraction and proteomic analysis

Colonoids were harvested in Cultrex Organoid Harvesting Solution and centrifuged at 300 xg for 10’ at 4°C. The cells were washed with ice cold PBS 5 times to remove any serum proteins. Cells were lysed in 250 μl of lysis buffer (60 mM HEPES pH 7.4, 150 mM KCl, 5 mM Na_3_EDTA, 5 mM EGTA, 1 mM Na_3_VO_4_, 50 mM NaF, 1 mM PMSF, 2% SDS (all MilliporeSigma)) supplemented with 1:100 of protease inhibitor cocktail (P8340, MilliporeSigma). Cells incubated with lysis buffer were sonicated on ice 3 times for 10 sec using 30% energy input. The lysed cells were centrifuged for 10 min at 5000 rpm at 4°C (MC2 Centrifuge, Sarstedt Desaga) to remove any unbroken cells. Protein concentration was determined by Bradford assay (Bio-Rad). Lysate was stored at −80°C. Proteomic analysis was carried out by the Mass Spectrometry and Proteomics Facility, Johns Hopkins University School of Medicine. Raw data was sent to and analyzed by Creative Proteomics (Shirley, NY). Figure S1A and B were generated by Creative Proteomics.

### RNA isolation and gene expression analysis

Colonoids were harvested from Matrigel using Cultrex Organoid Harvesting Solution. Cells were centrifuged at 5000 rpm for 5 min at 4°C. Supernatant was removed and pellet was stored at −80C until RNA extraction. RNA isolation was carried out using PureLink RNA Mini Kit (ThermoFisher) according to the manufacturer’s protocol. RNA concentration was determined using a DU 800 spectrophotometer (Beckman Coulter, Brea, CA). 500 ng to 2 ug of RNA was retro-transcribed into cDNA using SuperScript VILO Master Mix (ThermoFisher). DNA Real-time qPCR were run using PowerUp SYBR green Master Mix and QuantStudio 12K Flex Real-Time PCR instrument (all Applied Biosystems, Foster City, CA). Each sample was analyzed in triplicate. The primer oligonucleotide sequences are listed in Supplementary Table 3 (Xiaowei Wang, Athanasia Spandidos, Huajun Wang and Brian Seed: PrimerBank: a PCR primer database for quantitative gene expression analysis, 2012 update) AND (Bunaciu RP et al. 2008). The relative fold changes in mRNA levels between EspP-injured and control colonoids were determined using the 2^-∆∆CT^ method with normalization to *18S* ribosomal RNA.

### Statistics

Data are represented as means ± SEM. Statistical significances were calculated using Student’s *t*-test. Significance was represented as at least *p* < 0.05. All experiments were performed on a minimum of 3 different colonoid lines derived from separate normal human subjects, with a total of 7 colonoid lines used throughout these studies (Supplementary Table 1). N refers to number of independent replicates performed. All analyses were performed on GraphPad Prism 7.03 (GraphPad Software, La Jolla, CA).

---

**Figure S1.**
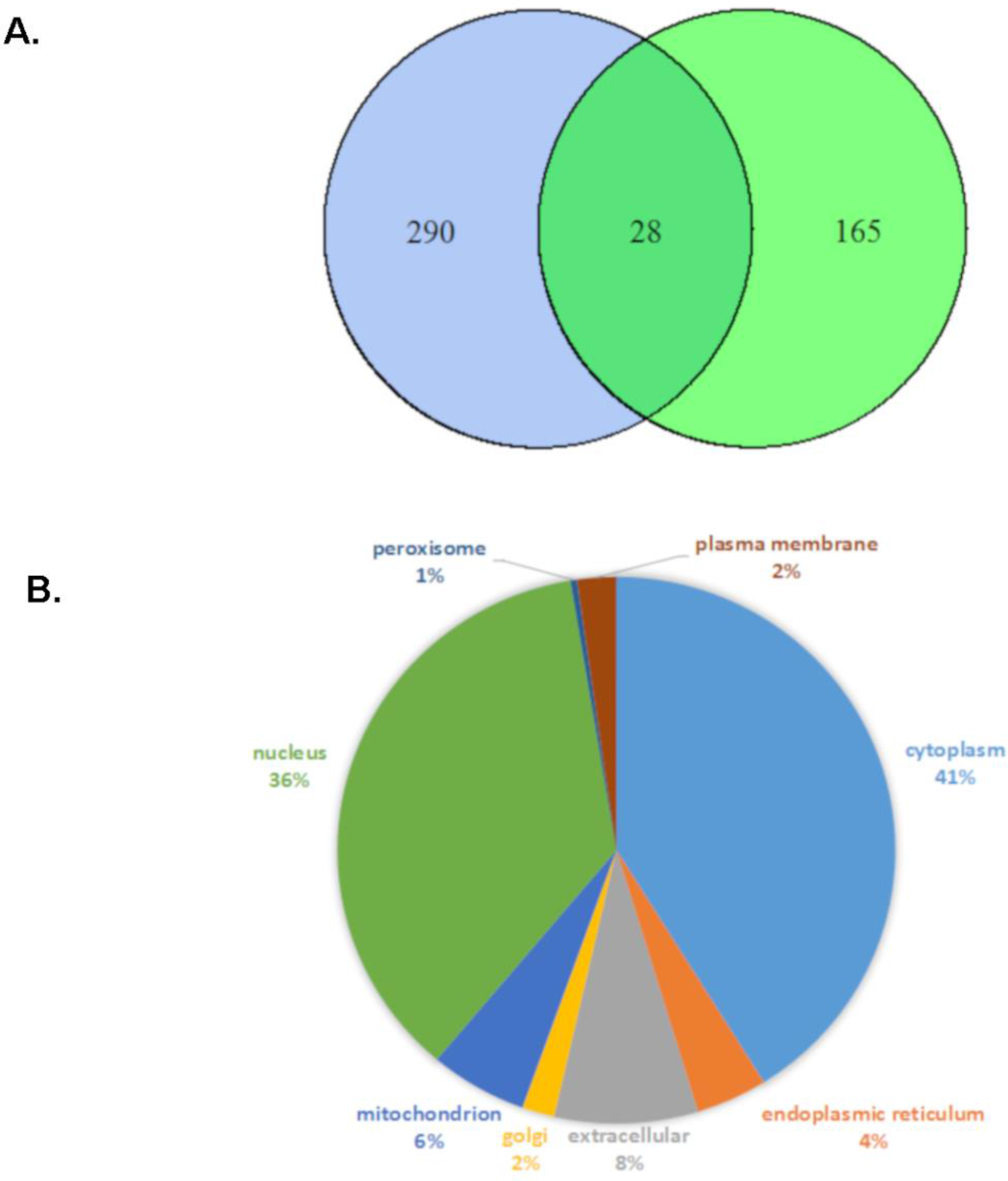
Proteomics analysis of differentially expressed proteins after EspP or EspP 263A treatment compared to control. (A) The Venn diagram depicts the number of differentially expressed proteins in the EspP S263A-treated (blue circle) and the EspP-treated (green circle) colonoids compared to control. Note the minimal overlap between the two treatments. (B) The distribution of subcellular localization of differentially expressed proteins in the EspP-treated compared to control colonoids.

**Figure S2.**
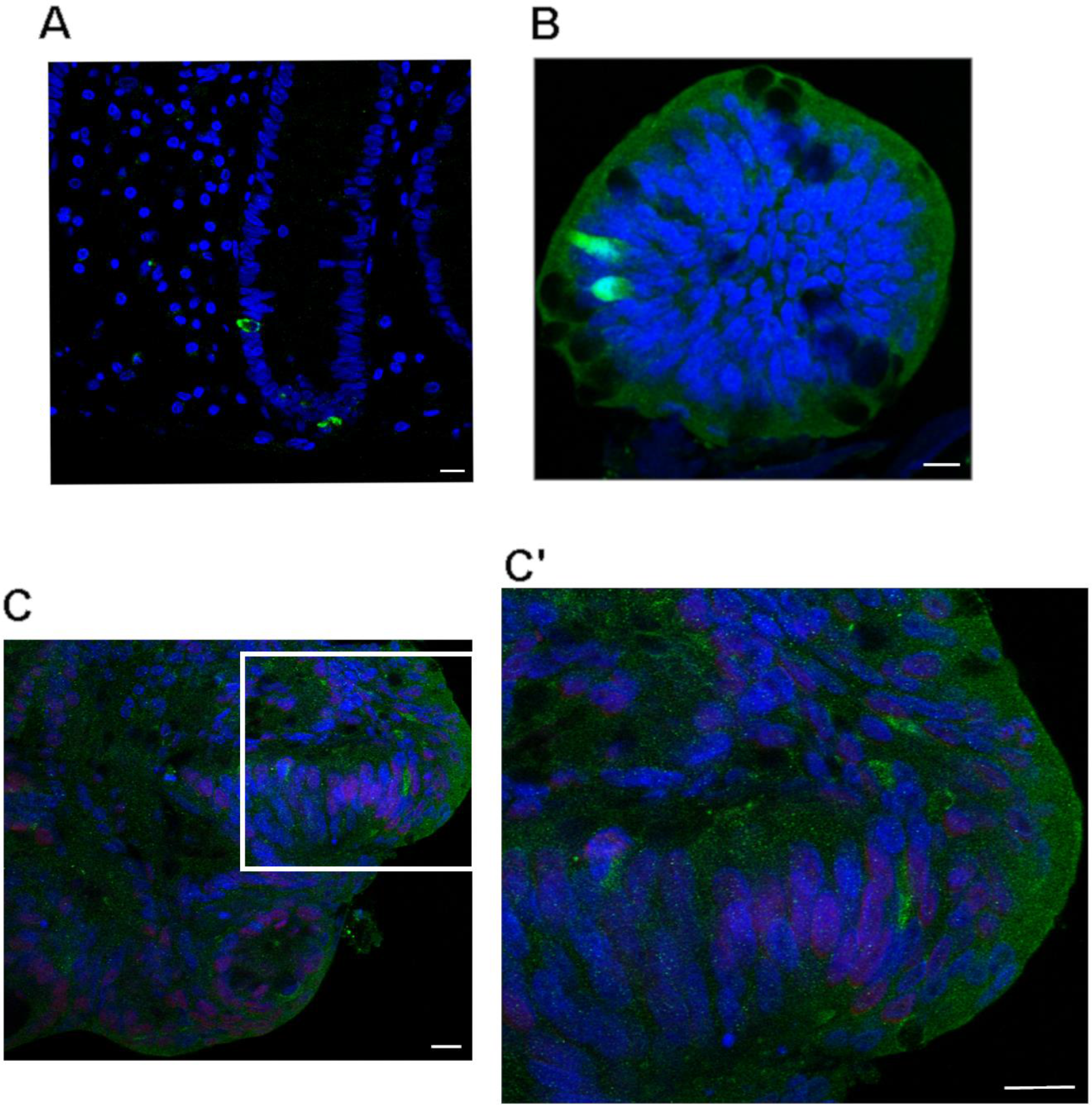
WNT2B marks a specific cell in the colonic crypt. (A and B) WNT2B is concentrated in a specific cell in (A) human colonic tissue and (B) human colonoids. WNT2B, green; nuclei, blue. (C) EspP-treated colonoids regenerating 24h post-EspP washout. WNT2B staining is more diffuse with more WNT2B+ cells, seen in the zoomed inset (C’); scale bar = 10 μm; WNT2B, green; nuclei, blue.

**Figure S3.**
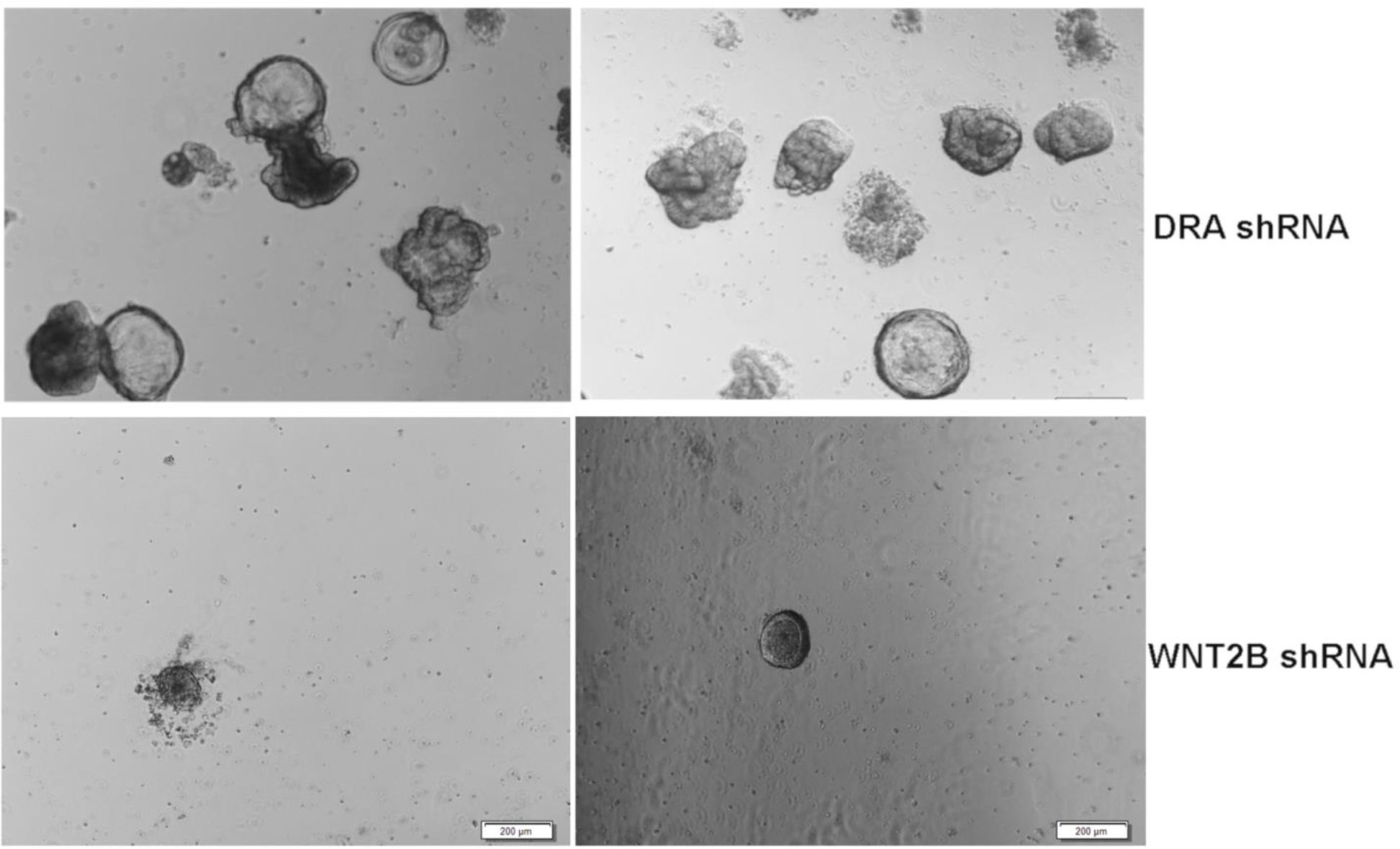
Knockdown of WNT2B results in non-viable colonoids. shRNA against DRA (top panel) and WNT2B (bottom panel) was introduced into duodenal enteroids or colonoids, respectively via lentivirus transduction. Images were taken 17 days post-transduction, showing healthy duodenal enteroids but lackluster colonoids.

**Supplemental Table 1.**
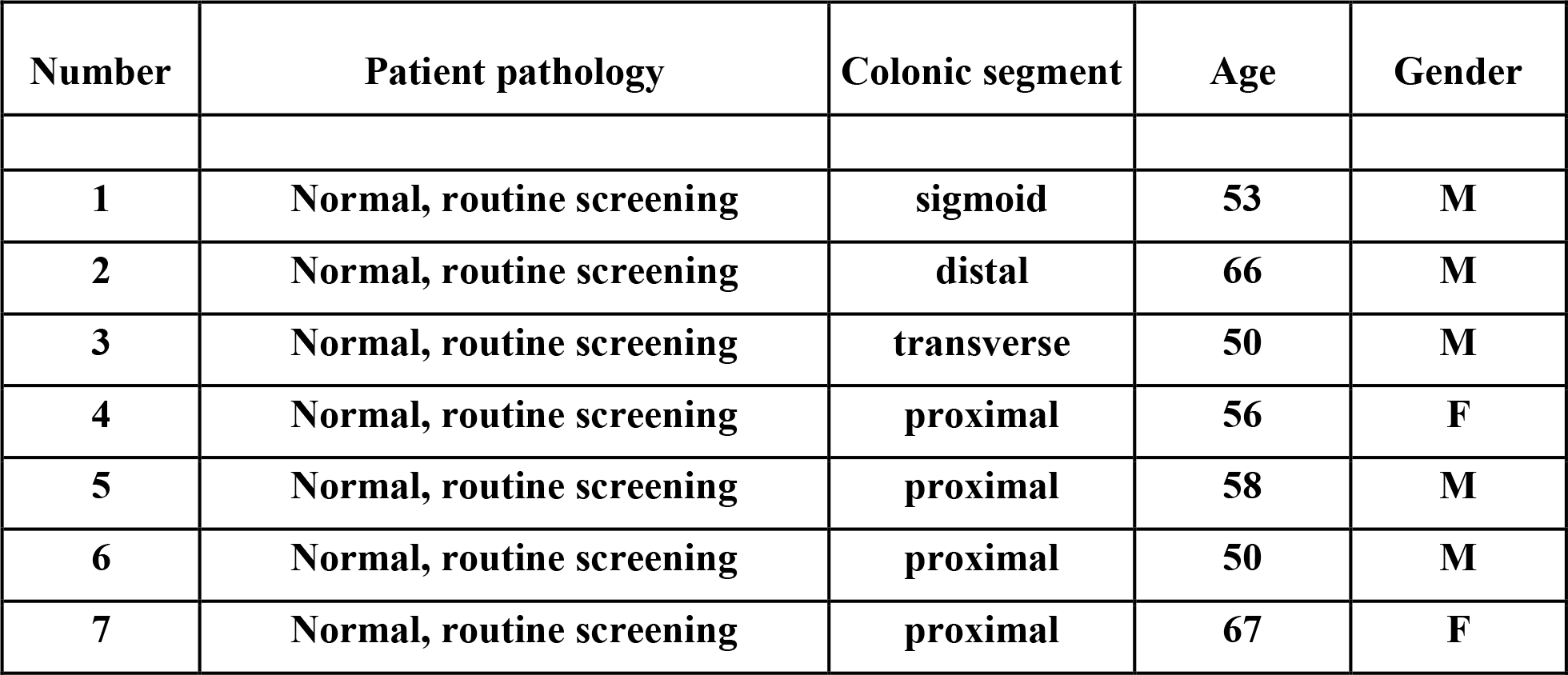

| **Number** | **Patient pathology** | **Colonic segment** | **Age** | **Gender** |
| --- | --- | --- | --- | --- |
| **1** | **Normal, routine screening** | **sigmoid** | **53** | **M** |
| **2** | **Normal, routine screening** | **distal** | **66** | **M** |
| **3** | **Normal, routine screening** | **transverse** | **50** | **M** |
| **4** | **Normal, routine screening** | **proximal** | **56** | **F** |
| **5** | **Normal, routine screening** | **proximal** | **58** | **M** |
| **6** | **Normal, routine screening** | **proximal** | **50** | **M** |
| **7** | **Normal, routine screening** | **proximal** | **67** | **F** |

**Supplemental Table 2.**
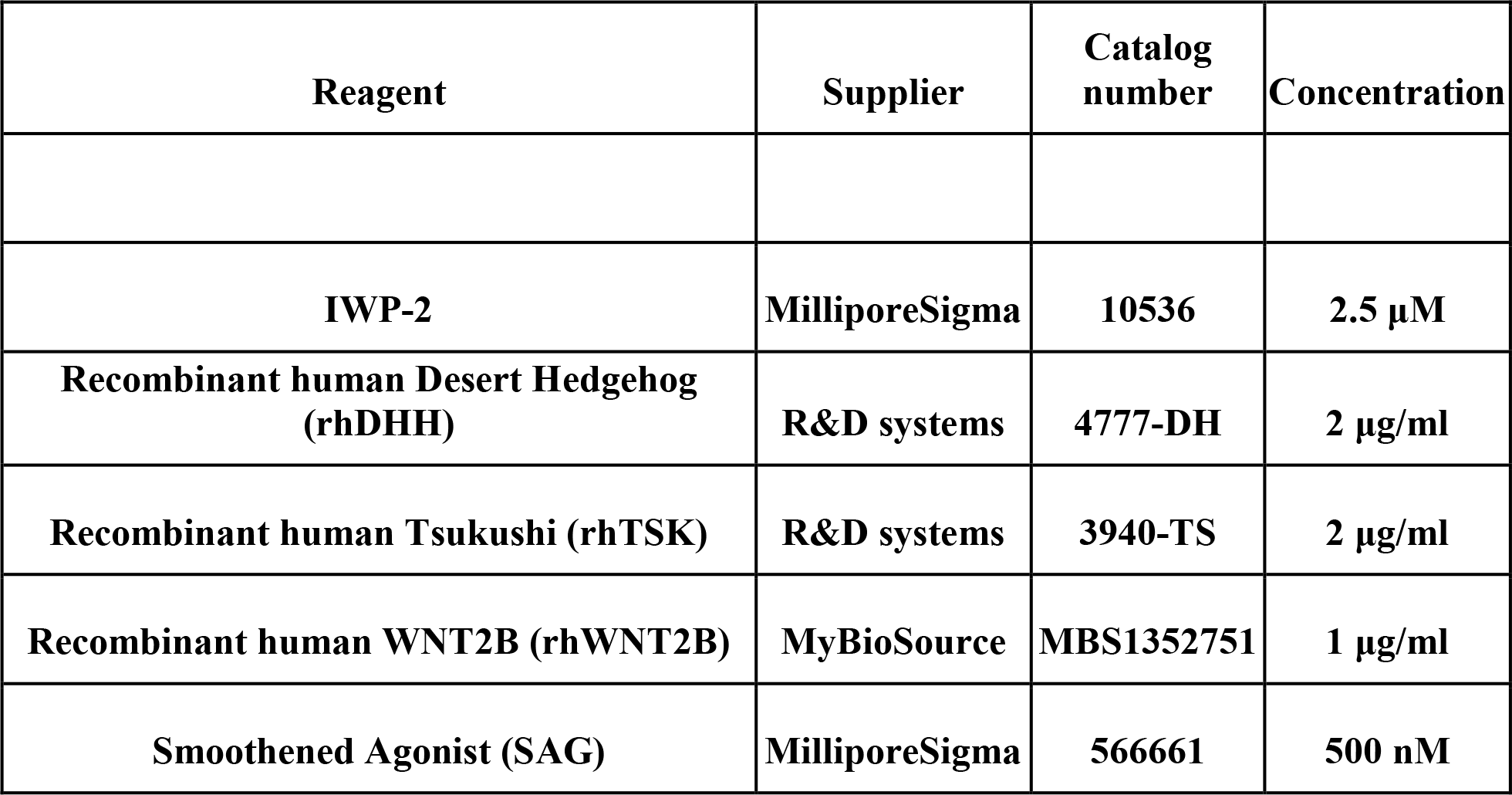

**Supplemental Table 3.**
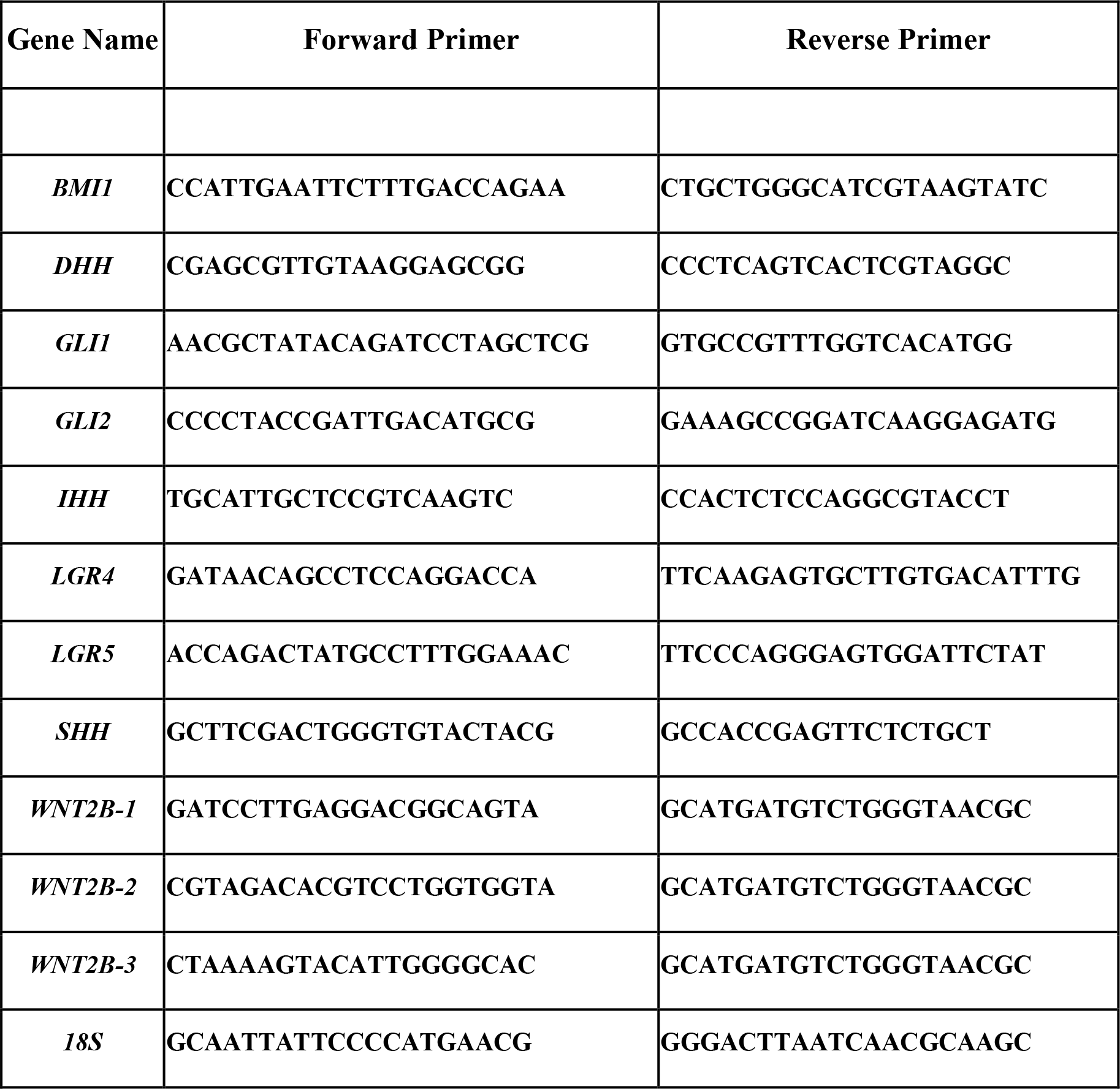

